# A bioprinting approach for high-throughput production of micropatterned neuroepithelial tissues and modeling TSC2-deficient brain malformations

**DOI:** 10.1101/2025.03.31.646426

**Authors:** Negin Imani Farahani, Kenneth Kin Lam Wong, George Allen, Abhimanyu Minhas, Lisa Lin, Shama Nazir, Lisa M Julian

**Author notes:** These authors contributed equally (Negin Imani Farahani, Kenneth Kin Lam Wong).

## Abstract

*In vitro* human pluripotent stem cell-derived models have been crucial in advancing our understanding of the mechanisms underlying neurodevelopment, though knowledge of the earliest stages of brain formation is lacking. Micropatterning of cell populations as they transition from pluripotency through the process of neurulation can produce self-assembled neuroepithelial tissues (NETs) with precise spatio-temporal control, enhancing the fidelity of hPSC models to the early developing human brain and their use in phenotypic assessments. Here, we introduce an accessible, customizable and scalable method to produce self-assembled NETs using bioprinting to rapidly deposit reproducibly sized extracellular matrix droplets. Matrix addition to the media provides a scaffold that promotes 3D tissue folding, reflecting neural tube development. We demonstrate that these scaffolded NETs (scNETs) exhibit key architectural and biological features of the human brain during normal and abnormal development, notably hyperproliferation and structural malformations induced by *TSC2*-deficiency, and provide a robust drug screening tool.

## Introduction

Given their high fidelity to human-specific neural cell types, and the inaccessibility of native brain tissue, *in vitro* neural cell and organoid models derived from human pluripotent stem cells (hPSCs) are transforming our understanding of the normal and disordered brain. However, the brain’s earliest stage–neurulation–through which the neuroepithelium forms and produces the neural tube, has been severely under-studied in humans. This is a critical time for healthy brain development as neural tube defects (~2 per 1000 pregnancies),^1^ which cause lethality or severe disabilities, and often malformations of cortical development (MCDs), a leading cause of refractory epilepsy and altered cognition,^2–4^ arise from neurodevelopmental disruptions at this early stage. Thus, the advancement of human models that permit detailed interrogation of neurulation-stage tissues is critical to uncover the mechanisms of onset for many neurodevelopmental disorders and to identify pathogenic mutations. Standard two-dimensional (2D) cultures do not suffice as they lack strict control over cell density, resulting in variable cell behaviors,^5^ and do not recapitulate morphogenic or architectural brain features. Three-dimensional (3D) organoids are also insufficient as they are inherently heterogeneous, exhibit abnormal architecture–producing multiple ventricular lumen–and are technically challenging to analyze during the first week of development when neurulation occurs.

Micropatterning- or microfluidic-based approaches to produce single lumen NETs from hiPSCs have gained popularity.^6–10^ The approach recapitulates structural aspects of the *in vivo* microenvironment by restricting the cell growth surface during neural lineage induction (neurulation), which facilitates self-assembly of newly formed neural stem cells into neuroepithelial tissues that surround a single lumen. These tissues reflect the *in vivo* architecture of the neural tube with physiologically relevant metrics of cell and architectural complexity. Current approaches to generate micropatterned tissues include costly, laborious and often low-throughput technologies like photolithography, soft lithography, surface passivation and contact photo patterning which, although effective, often require specific expertise.^11^

Here we establish a customizable, accessible and scalable method to produce neuroepithelial micropatterned tissues using a bioprinter to generate reproducibly sized matrix droplets for hPSC adhesion prior to neural induction. Inclusion of Matrigel in the culture media provides a scaffold for 3D tissue folding.^8^ We demonstrate that these scaffolded neuroepithelial tissues (scNETs) are a reproducible physiologically relevant model of early neurodevelopment. We also show that scNETs display striking hyperproliferation and tissue malformations induced by mutations in the mTORC1 inhibitor TSC2, which cause the MCD disorder tuberous sclerosis (TS). Thus, scNETs offer a strikingly reproducible and quantifiable model of early MCD formation, and we demonstrate their potential for high-throughput phenotyping and drug screening applications.

## Results

### Generation of hPSC-derived scaffolded neuroepithelial tissues (scNETs) by micropatterning on bioprinted matrices

We set out to develop a scalable physiologically relevant model of early brain development by harnessing hPSCs, neural lineage induction, and bioprinting technologies to generate micropatterned scNETs. We used an extrusion bioprinter to print individual droplets of the extracellular matrix (ECM) basement membrane formulation Cultrex, up to 100 droplets in 3 minutes, with a defined diameter onto the surface of cell culture vessels **(Figure 1A)**. This enabled spatially restricted growth of hPSCs seeded onto the ECM droplets. We could reliably print droplets as small as 500 µm in diameter **(Figure S1A)** though 800 µm droplets were used for most experiments, which corresponds to the diameter of the Carnegie stage 13-14 (4-5 gestational weeks) human forebrain neural tube.^12,13^ Highlighting the customizability of our approach, we used G-code, a programming language for 3D bioprinting, to instruct the printer to deposit arrays of droplets with reproducible size and precise spatial arrangements **(Figure S1B-C)**. This allows for easy modification of the number, size, shape, and location of droplets to meet the requirements of each experiment. Droplets can be printed on various types of tissue culture ware, including petri dishes, multi-well plates and coverslips **(Figure S1D)**, demonstrating the flexibility, scalability and high throughput compatibility of our method.

**Figure 1.**
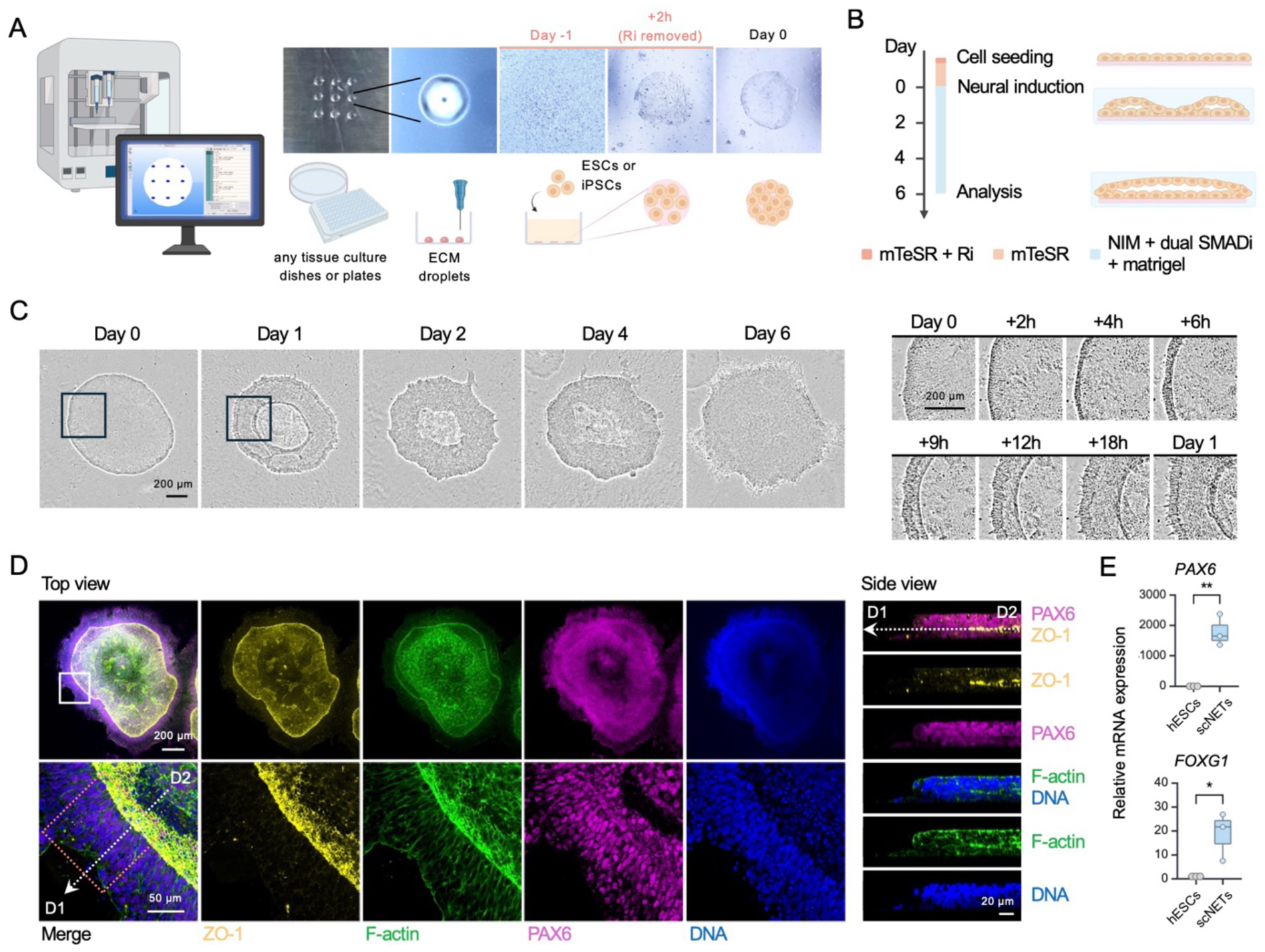
A programmable bioprinting approach to generate hPSC-derived, self-organized, micropatterned scNETs. **(A)** Left to right: Schematic of a 3D bioprinter and G-code used to deposit ECM droplets onto the surface of a culture dish or multi-well plate. Next, an image of 9 ECM droplets; magnified view of a droplet; hPSC seeding the day before neural induction (Day −1); removal of ROCK inhibitor (Ri) and unattached cells 2 hours (h) post-seeding; formation of a circular hPSC colony. **(B)** Schematic of the timeline and media used to generate micropatterned scNETs (left) and side views of monolayered hPSCs, then developing scNETs following neural induction. **(C)** Phase-contrast time-lapse imaging of scNET development starting from the day of neural induction to day 6 (left). Magnified views of the boxed regions showing thickening (multilayering) at the edge of scNETs (right). See also **Video 1**. **(D)** Confocal images of top and side views of a day 6 scNET (H1) stained for ZO-1 (yellow), F-actin (green), PAX6 (magenta) and DAPI/ DNA (blue). Top view: the bottom row shows magnified views of the white boxed inset drawn on the first image in the top row. **Figure 2A** shows the magnified views of the dashed red inset. The dashed arrow D1-D2 refers to the *z-*section shown in the side view images. See also **Video 2**. **(E)** Boxplot showing relative mRNA levels of *PAX6* and *FOXG1* in undifferentiated hESCs (H1) and day 6 scNETs. *n* = 3 biological replicates. \*\**p* < 0.005, \**p* < 0.05, unpaired *t-*test. Scale bars 200 µm, 50 µm and 20 µm are shown as indicated. See also **Figures S1 and S2**. Next, we asked if self-organized neural tissues could be produced from hPSCs seeded onto bioprinted matrix droplets by inducing neural differentiation from hPSCs by dual SMAD inhibition.^14,15^ After seeding, hPSCs attached to and proliferated within the area of printed ECM, forming monolayered, circular colonies **(Figure 1A)**. Once hPSCs reached confluence neural differentiation was initiated **(Figure 1B)**, marking day 0 of the assay. We included 4% Matrigel in the medium which, as previously reported,^8^ should provide a scaffold for neural folds to project and converge, driving a transition from 2D culture into 3D single lumen neural tube-like tissues.

**Figure S1.**
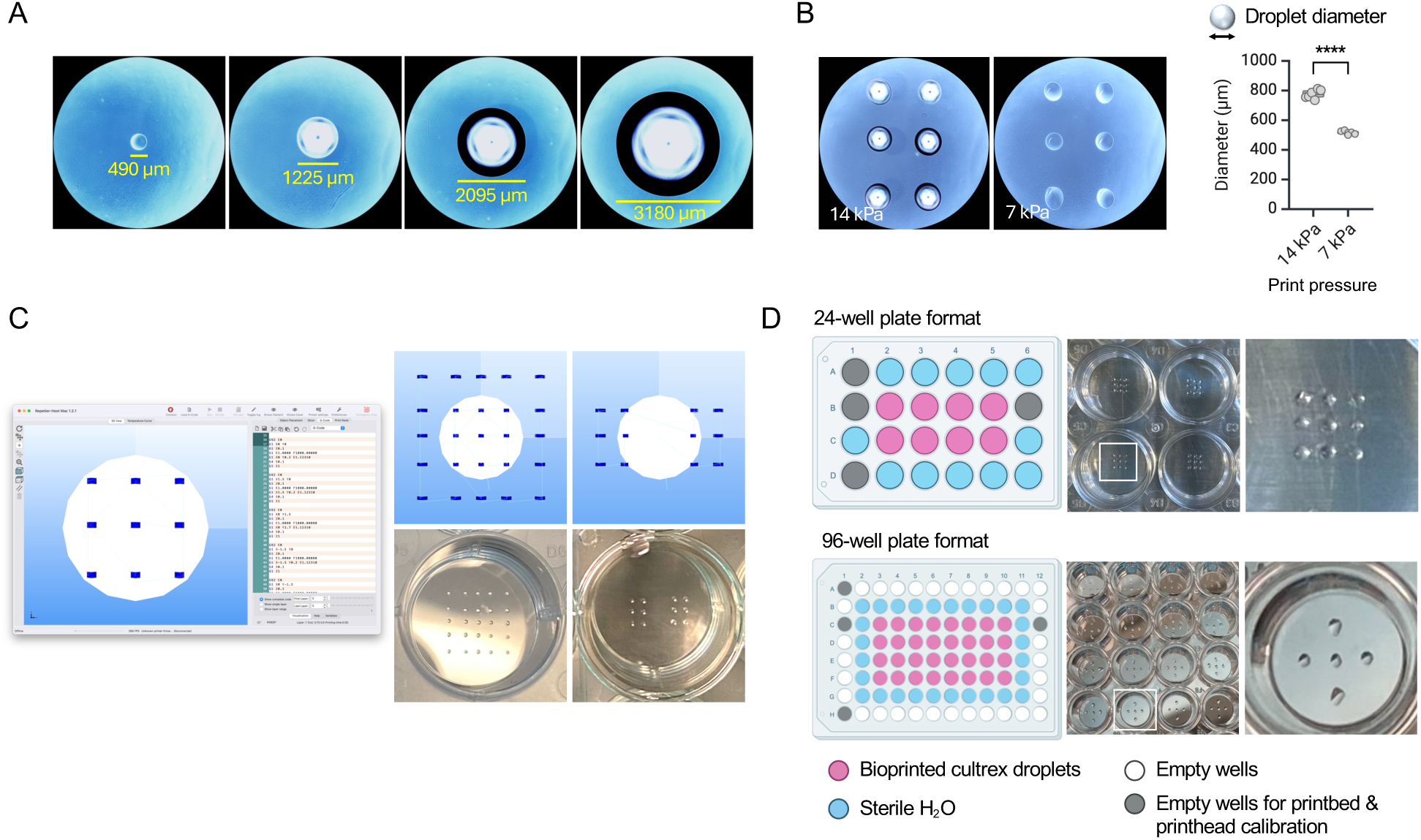
Programmable bioprinting of geometrically-defined ECM droplets at assigned positions. **(A)** Brightfield images showing bioprinted ECM droplets of indicated diameters. **(B)** Brightfield images showing bioprinted ECM droplets using printhead pressure at either 14 or 7 kPa (left). Boxplot showing the droplet diameter at the indicated pressure. Note that we were able to consistently print arrays of ECM droplets of a similar size. \*\*\*\**p* < 0.00005, unpaired *t-*test. **(C)** Screenshots of Repetier-Host showing programmable printing arrangements using G-code (left). More examples showing diverse droplet arrangements printed (right). **(D)** Bioprinting in 24-well (left) and 96-well (right) plate formats, with photos and zoom-in views showing the droplets. Wells in grey were used for print bed and printhead calibrations. Wells in blue were filled with sterile water to minimize droplet evaporation. Wells in red were bioprinted with ECM droplets. Wells in both grey and white were empty.

Within 2-4 hours of neural induction multilayering occurred, indicated by the thickening of the micropatterns starting from the edge and progressing inward, and by the end of day 1 a ring-shaped structure had emerged **(Figure 1C**, **Figure S2A, Video 1)**. The accumulation of filamentous actin (F-actin) between cell layers suggests early spatial organization. This self-organizing process was reproducible across different hPSC lines, including previously described human embryonic stem cells (hESCs: H1) and induced pluripotent stem cells (iPSCs: 168) **(Figure S2B).**^16,17^ The micropatterned tissues continued to grow and thicken, ultimately forming a circular disc-like structure by day 6 **(Figure 1D, Video 2)**. They also developed a central lumen or ventricle-like structure, with its borders defined by enriched ZO-1 (a tight junction scaffolding protein) and F-actin (**Figure 1D**). Notably, most cells stained positive for PAX6, confirming their transition into neural stem and progenitor cells (NPCs) with a dorsal forebrain identity. This was further corroborated by RT-qPCR analysis, which showed significantly higher expression of *PAX6* and *FOXG1*, another dorsal telencephalon marker, in scNETs compared to undifferentiated hPSCs **(Figure 1E**, **Figure S2C)**.

**Figure S2.**
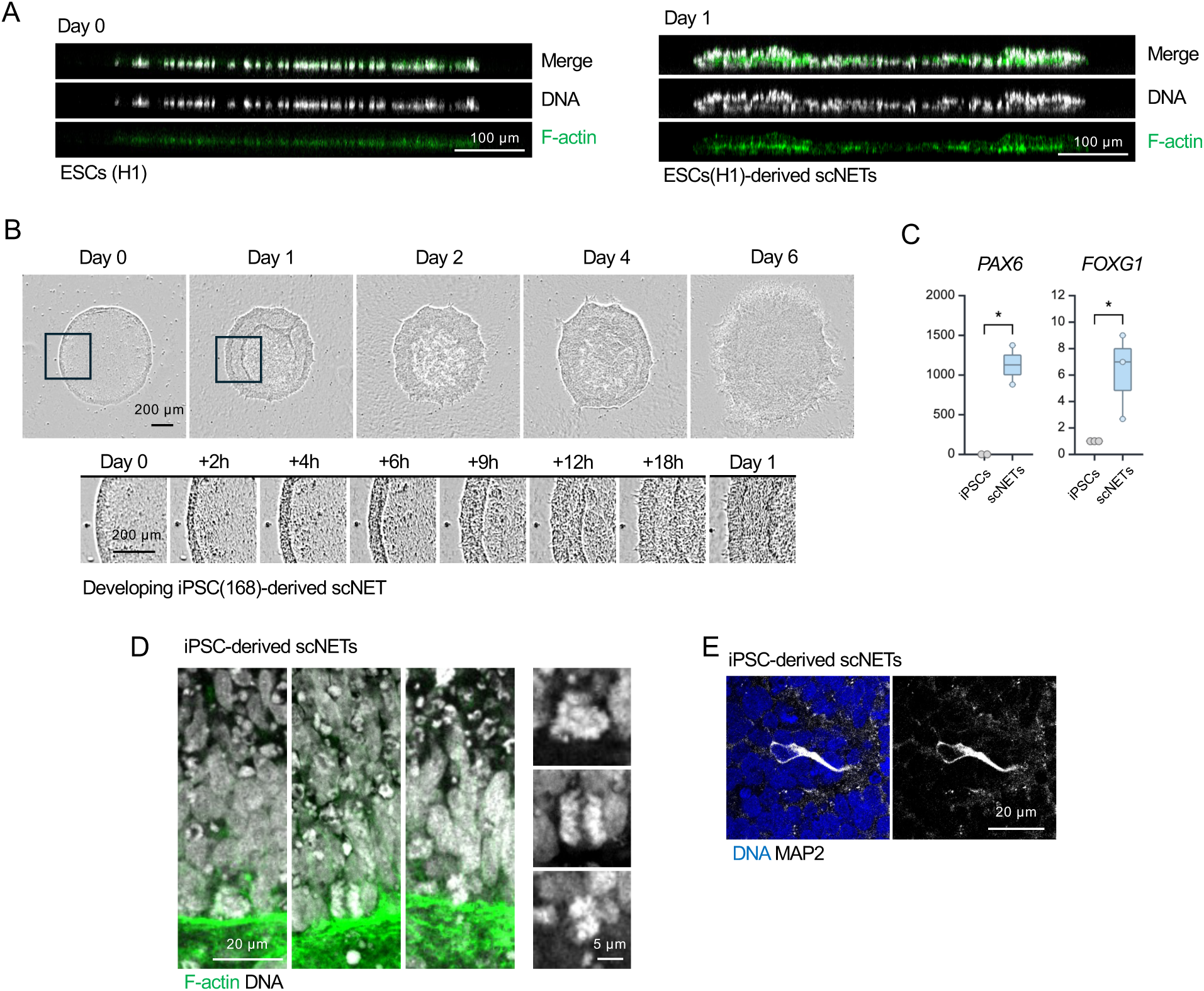
Developing scNETs self-assemble into 3D neuroepithelial tissue-like structure with neurogenic potential. **(A)** Confocal images of undifferentiated hESCs (H1) on day 0 and developing scNETs (H1) on day 1, stained for F-actin (green) and DNA (grey). **(B)** Phase-contrast time-lapse imaging of the development of scNET derived from iPSCs (168) starting from day 0 till day 6 (left). Zoom-in views of the insets showing the edge thickening that took place between day 0 and day 1. Scale bars indicate 200 µm. Refer to **Video 1**. **(C)** Boxplot showing relative mRNA levels of *PAX6* and *FOXG1* of day 6 scNETs and the undifferentiated iPSCs (168). \**p* < 0.05, unpaired *t-*test. **(D)** Confocal images of the neuroepithelium-like layer of scNETs derived from iPSCs (168), stained for F-actin (green) and DNA (blue). Scale bars 20 µm and 5 µm are shown. **(D)** Confocal images of a MAP2^+^ cell in scNETs derived from iPSCs (168), stained for MAP2 (grey) and DNA (blue). Scale bar indicates 20 µm.

### scNETs are a physiologically relevant model for early neurodevelopment

We further investigated the relevance of the scNET model to human neural tube development. As the neural tube forms *in vivo*, neuroepithelial cells become polarized with the apical domain adjacent to the ventricular lumen and the basal domain extended towards the outer surface of the neural tube^18^ This apicobasal polarity is crucial for proper interkinetic nuclear migration (IKNM), a process by which the nuclei of NPCs migrate toward the basal surface of the neuroepithelium during G1 to S phases, then back to the apical surface for mitosis.^19^ IKNM plays a vital role in the integrity of cell cycle progression and the fate decisions NPCs make.^20,21^

ZO-1 and F-actin enrichment demarcates the scNET lumen **(Figure 1D, 2A)**. Since ZO-1 marks the apical surface in polarized epithelia,^22^ its localization in the tissues confirmed that PAX6^+^ NPCs had acquired apicobasal polarity, with their apical side facing the center. Enrichment of apical marker N-cadherin at the lumen side **(Figure 2B)** further highlights apicobasal polarity. We also observed that though some apical NPCs appeared circular (arrowhead **Figure 2A**, **Figure 2C2)**, most were elongated and columnar. This cellular arrangement resembles the pseudostratified neuroepithelium of the developing forebrain, in which NPCs appear multilayered due to the positioning of their nuclei at different heights.^23,24^

**Figure 2.**
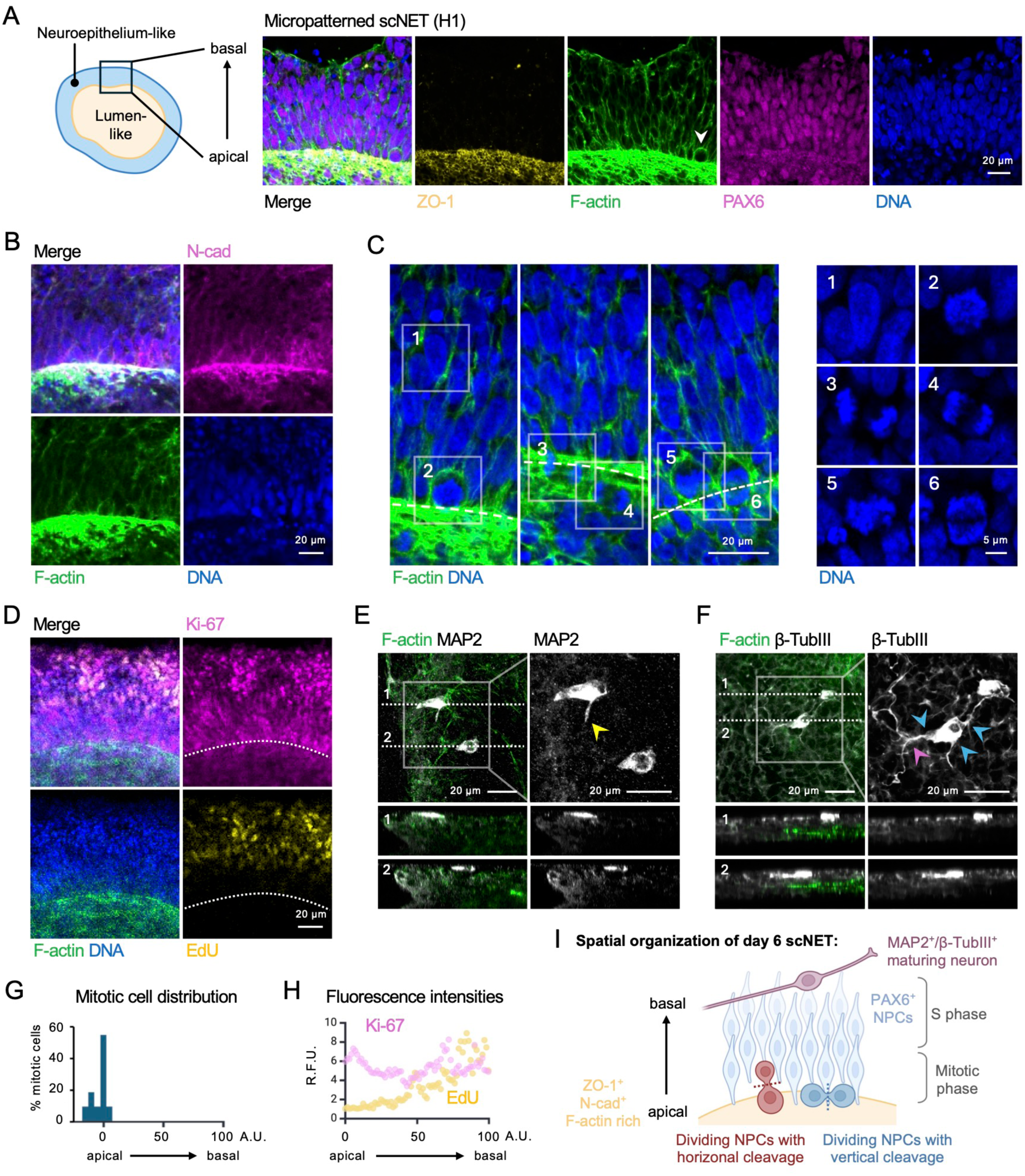
Micropatterned scNETs closely resemble the early neuroepithelium and produce the first neuronal cells within 6 days. **(A)** Schematic of a day 6 scNET highlighting the peripheral neuroepithelium and central lumen structures (left). Magnified views of the dashed red inset drawn in **Figure 1D**, showing the neuroepithelium-like layer oriented with apicobasal polarity (right); ZO-1 (yellow), F-actin (green), PAX6 (magenta) and DAPI/ DNA (blue). Arrowhead marks a circular PAX6^+^ NPC. **(B)** Confocal images of a day 6 scNETs (H1) stained for N-cad (magenta), F-actin (green), and DNA (blue). **(C)** Confocal images showing the distribution of non-dividing and dividing cells in day 6 scNETs (H1), stained for F-actin (green) and DNA (blue) (left). Magnified views of the insets (1-6) (right). Dashed lines mark the apical surface. **(D)** Confocal images of a day 6 scNET (H1) stained for Ki-67 (magenta), EdU (yellow), F-actin (green), and DNA (blue). Dashed lines mark the apical surface. **(E-F)** Confocal images showing a scNET (H1) stained for MAP2 (grey) and F-actin (green) **(E)**, and a scNET (168) stained for β-TubIII (grey) and F-actin (green) **(F)**. Yellow arrowhead marks a developing neurite. Blue arrowheads mark multiple neurites from the same cell. Purple arrowheads point to the branching of the neurite. Dashed lines refer to the region at which side view images were taken. **(G)** A distribution graph showing the percentage of mitotic cells across the apicobasal axis of the neuroepithelium layer of scNETs (H1), using apicobasal distance scaled from 0-100 arbitrary units (A.U.), with apical surface marked by 0, basal surface 100, and luminal region negative. **(H)** A scatter plot showing the relative fluorescence units (R.F.U.) of Ki-67 (purple) and EdU (yellow) across the apicobasal axis of the neuroepithelium layer of scNETs (H1), using apicobasal distance scaled from 0-100 arbitrary units (A.U.), with apical surface marked by 0 and basal surface 100. **(I)** Schematic showing the spatial organization of PAX6^+^ NPCs and MAP2^+^/β-TubIII^+^ maturing neurons in scNETs generated through our bioprinting approach. Scale bars 20 µm and 5 µm are shown as indicated. See also **Figure S2**.

The presence of circular NPCs at the apical surface prompted us to examine the distribution of mitotic cells along the ventricle, as DAPI staining revealed these cells had condensed DNA **(Figure 2C2-6)**. Whereas non-dividing cells displayed diffuse DNA **(Figure 2C1)**, the condensed DNA of the dividing cells signified entry into prophase **(Figure 2C2, 2C5)**. We also identified cells in anaphase, denoted by the separation of sister chromatids towards opposite ends of the cell **(Figure 2C3-4, 2C6)**, with cleavage planes either perpendicular or parallel to the apical surface (**Figure 2I**). Notably, mitotic cells were predominantly localized at the apical surface **(Figure 2C, 2G)**, as observed in the native neuroepithelium.^23,21^

To assess NPC proliferation states, we examined the expression of Ki-67, a nuclear protein that accumulates in proliferating cells and is degraded in quiescence.^25^ Immunostaining showed that Ki-67 was highly expressed in scNETs at both the apical and basal regions **(Figure 2D, 2H)**. 5-ethynyl-2′-deoxyuridine (EdU) labeling, which marks cells replicating their DNA, showed that cells in the DNA synthesis phase of the cell cycle were enriched toward the basal side **(Figure 2D, 2H)**. This spatial distribution of mitotic cells at the apical surface and S-phase cells at the basal side is characteristic of the natural IKNM process.

The orientation of the mitotic cleavage plane is known to infer whether a neural stem cell is undergoing a symmetric self-renewing division or an asymmetric differentiative division.^21,23,26^ While most mitotic divisions had a cleavage plane parallel or vertical to the apical surface, suggesting proliferative self-renewing divisions, cells with a horizontal cleavage plane were observed, suggesting a low level of asymmetric divisions that might generate early neurons (**Figure 2C**). Immunostaining for β-Tubulin III and MAP2 confirmed the presence of immature and maturing neurons by day 6 in the scNETs **(Figure 2E-F**, **Figure S2E)**. These cells were localized at the basal side, characteristic of developing neurons *in vivo*.^27^

Together, these findings demonstrate that PAX6^+^ NPCs within scNETs recapitulate key features of the neuroepithelium including apicobasal polarity, pseudostratification, IKNM and early stages of neurogenesis (**Figure 2I**), making this a highly relevant model to study early corticogenesis.

### scNETs generated from *TSC2^−/−^* hPSCs display hyperproliferation and cortical folding

The physiological relevance of scNETs prompted us to ask if disease-associated phenotypes could be identified and quantified in these tissues. We focused on TSC2, which acts in complex with TSC1 to inhibit the nutrient-sensing mammalian target of rapamycin complex 1 (mTORC1) signaling pathway.^28,29^ Inactivating mutations in the *TSC2* gene results in MCDs due to excess cell proliferation and altered differentiation, largely due to mTORC1 hyperactivation.^4,29–33^ Thus, we generated scNETs using *TSC2-*deficient hPSCs (*TSC2^−/−^)* that were previously engineered by CRISPR-Cas9 genome editing.^16,17^ We first confirmed the knockout mutation in these cells via PCR **(Figure S3A)**. One day after neural induction, the developing *TSC2^−/−^* scNETs were indistinguishable from wild-type (WT), characterized by the formation of a standard ring-like structure **(Figure 3A**, versus control in **Figure 1C)**. However, as early as day 2, unlike WT, *TSC2^−/−^*scNETs began to develop a scalloped border that is suggestive of cortical folding. Excess cortical folds or gyri have been observed in TSC2-driven MCDs including TS and polymicrogyria,^33–35^ though this phenotype has not been modeled in human organoids. These convex folds became more prominent as the *TSC2^−/−^* scNETs continued to grow **(Figure 3A)** and were observed in both H1 ESC- and 168 iPSC-derived scNETs **(Figure S3B-C)**.

**Figure 3.**
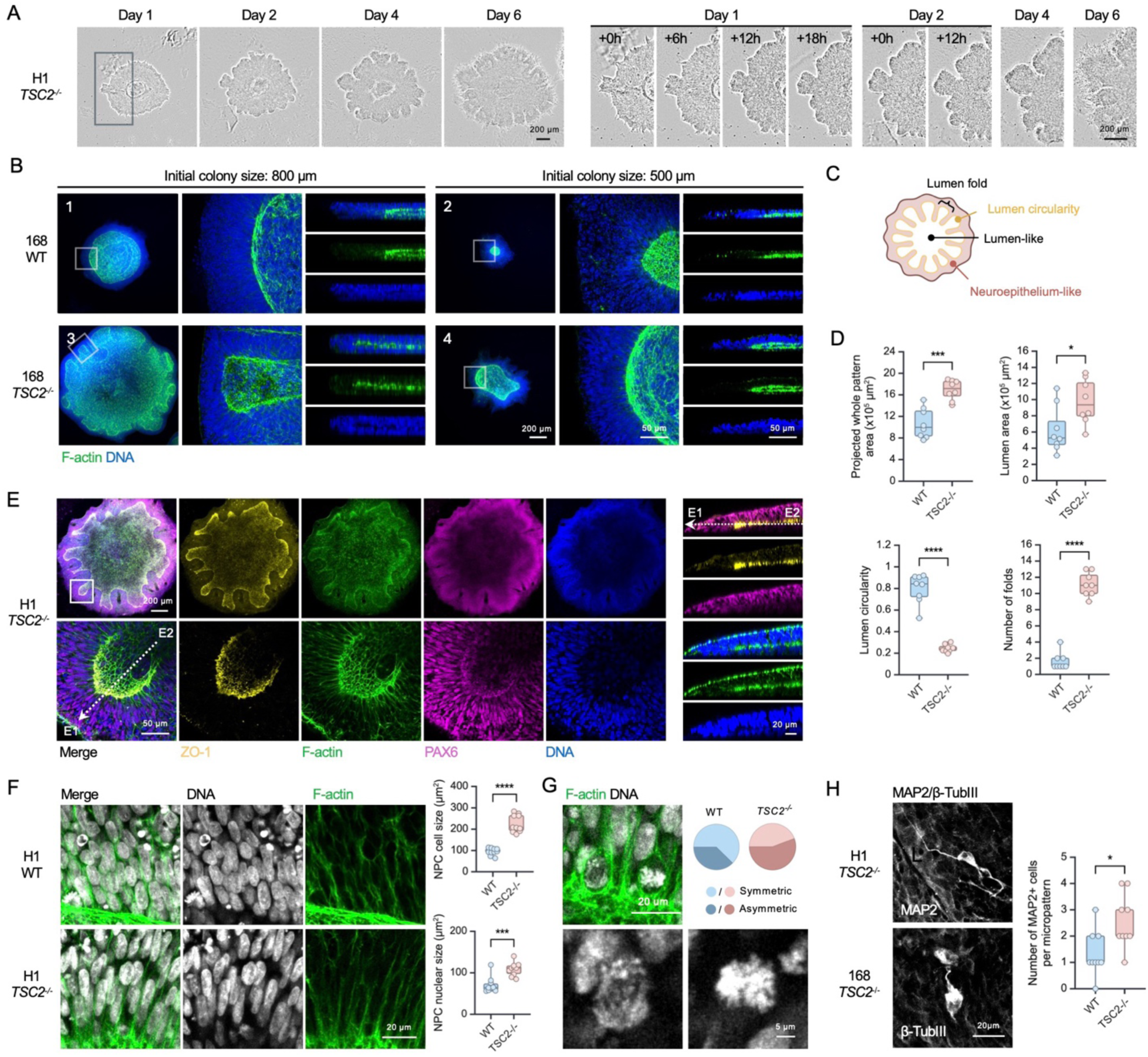
Micropatterned scNETs derived from *TSC2^−/−^* hPSCs display hyperproliferation and cortical folding phenotypes. **(A)** Phase-contrast time-lapse imaging of *TSC2^−/−^* scNET development (H1) (left). Magnified views of the inset showing the emergence of a cortical folding-like phenotype between day 1 and day 2. See also **Video 3**. **(B)** Confocal images of day 6 wildtype **(B1-B2)** and *TSC2^−/−^* **(B3-B4)** scNETs (168) with their indicated initial colony sizes stained for F-actin (green) and DNA (blue). **(C)** Schematic of the *TSC2^−/−^* scNET highlighting morphological abnormalities due to the loss of *TSC2* gene expression. **(D)** Boxplots showing projected whole pattern area, lumen area, lumen circularity and number of folds of day 6 WT scNETs (H1), compared with the *TSC2^−/−^* scNETs. \*\*\*\**p* < 0.00005, \*\*\**p* < 0.0005, \**p* < 0.05, unpaired *t-*test. **(E)** Confocal images of a day 6 *TSC2^−/−^* scNET (H1) stained for ZO-1 (yellow), F-actin (green), PAX6 (magenta) and DNA (blue). See also **Video 2**. **(F)** Confocal images showing the neuroepithelium-like layers of day 6 WT and *TSC2^−/−^* scNETs (H1), stained for F-actin (green) and DNA (grey). Boxplots showing the quantifications of cell size (outlined by F-actin) and nuclear size. \*\*\*\**p* < 0.00005, \*\*\**p* < 0.0005, unpaired *t-*test. **(G)** Confocal images showing mitotic cells of day 6 *TSC2^−/−^* scNETs (H1), stained for F-actin (green) and DNA (grey). Pie chart showing the percentage of symmetric and asymmetric divisions. **(H)** Confocal images showing MAP2^+^ and β-TubIII^+^ cells of day 6 *TSC2^−/−^* scNETs. Boxplot showing the number of MAP2^+^ cells in WT scNETs and that in *TSC2^−/−^* scNETs (H1). \**p* < 0.05, unpaired *t-*test. Scale bars 200 µm, 50 µm, and 20 µm are shown. See also **Figure S3**.

Taking advantage of the flexibility of our bioprinting approach to modify ECM droplet size, we asked if *TSC2^−/−^* phenotypes would be affected by initial colony size. WT scNETs grown on 500 µm ECM droplets were smaller than those grown on 800 μm droplets **(Figure 3B1-2)**. Both featured a central lumen structure. Interestingly, whereas *TSC2^−/−^* scNETs grown on 800 µm ECM islands displayed extensive folding, those on 500 μm developed fewer folds, indicated by the lumen circularity **(Figure 3B3-4)**. Yet, the *TSC2^−/−^*scNETs grown on 500 μm ECM droplets were still larger than their control counterparts **(Figure 3B2, 3B4)**. These data suggest the *TSC2^−/−^* hyperproliferative phenotype occurs regardless of the initial colony size but the folding phenotype occurs in larger *TSC2^−/−^* scNETs, possibly because a larger colony size may provide more room for the cells to grow and interact with one another.

PAX6^+^ NPCs were abundant in *TSC2^−/−^* scNETs, like WT, and displayed apicobasal polarity with the apical side marked by ZO-1 and differentiated into MAP2^+^/β-TubIII^+^ cells with visible neurites **(Figure 3E-H)**. We noted a significant increase in cell size in *TSC ^−/−^* scNETs **(Figure 3F)**, in line with previous reports.^36,37^ Our analysis further revealed increased asymmetric division in *TSC2^−/−^*compared to WT scNETs **(Figure 3G)**, supporting previous reports that TSC2 loss alters cell differentiation^38^ and that during early development this manifests as accelerated neurogenesis.^39^ Indeed, even at this very early stage of neurodevelopment, scNETs showed a slight increase in the number of MAP2^+^ cells **(Figure 3H)**. Altogether, we demonstrate that our bioprinting approach permits the generation of reproducible scNETs and the manifestation of MCD disease-associated phenotypes in just a few days.

**Figure S3.**
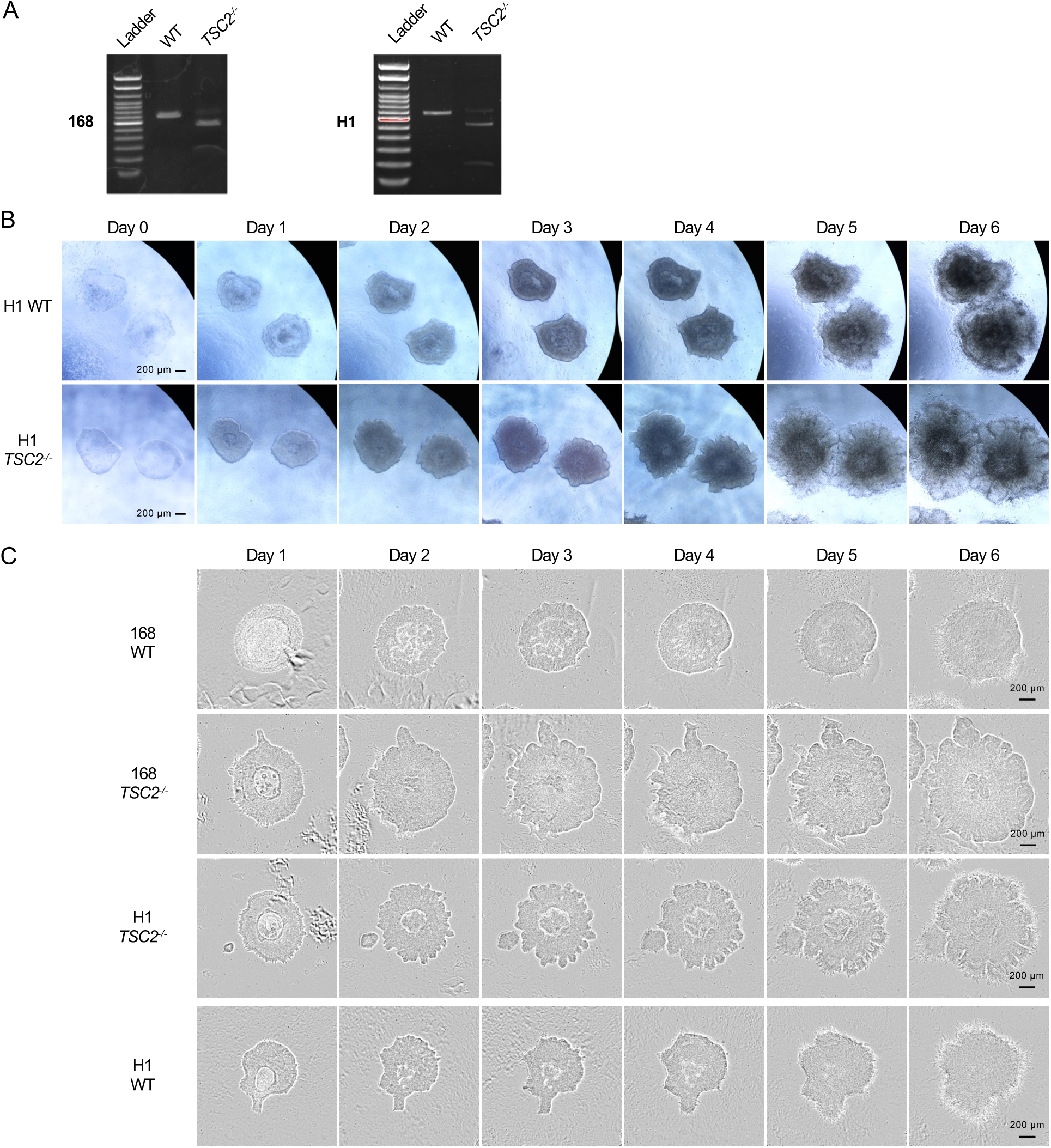
*TSC2^−/−^* scNETs display cortical folding-like phenotype as early as on day 2. **(A)** Verification of *TSC2^−/−^* cell lines (hESC H1 and iPSC 168) by PCR. See method section for details. **(B)** Brightfield images of developing scNETs derived from WT hESCs (H1) (top row) and those derived from *TSC2^−/−^* hESCs (H1). **(C)** Phase-contrast time-lapse imaging of developing scNETs derived from WT iPSCs (168) (1st row), *TSC2^−/−^* iPSCs (168) (2nd row), WT hESCs (H1) (3rd row) and *TSC2^−/−^* hESCs (H1) (4th row). Note that even though the initial shape of the WT scNET (H1) was irregular, it did not cause folding as extensive as the knockout counterpart.

### Rapamycin rescues *TSC2^−/−^* phenotypes in scNETs

Finally, we assessed the sensitivity of the scNET model to reflect phenotypic changes in response to drug treatment. Using levels of phosphorylated ribosomal protein S6 at Ser235/236 (pS6) as an indicator of mTORC1 activity,^16,40^ we confirmed mTORC1 hyperactivity in *TSC2^−/−^* scNETs compared to WT **(Figure 4B**). Treatment of scNETs with rapamycin (20 nM), an mTORC1 inhibitor that represents the leading clinical treatment of TS patients,^41–43^ for 48 hours starting on day 1 **(Figure 4A)** reduced pS6 levels **(Figure 4B**, **Figure S4D)**. Additionally, *TSC2^−/−^*scNETs treated with rapamycin (20 nM) were significantly smaller in size **(Figure 4C-D**, **Figure S4A-B)** than untreated *TSC2^−/−^* scNETs and had significantly fewer folds. KI67 immunostaining and EdU incorporation further showed that rapamycin rescued the increased proliferation observed in untreated *TSC2^−/−^* scNETs, as both were significantly decreased in treated tissues **(Figure 4C-D)**.

**Figure 4.**
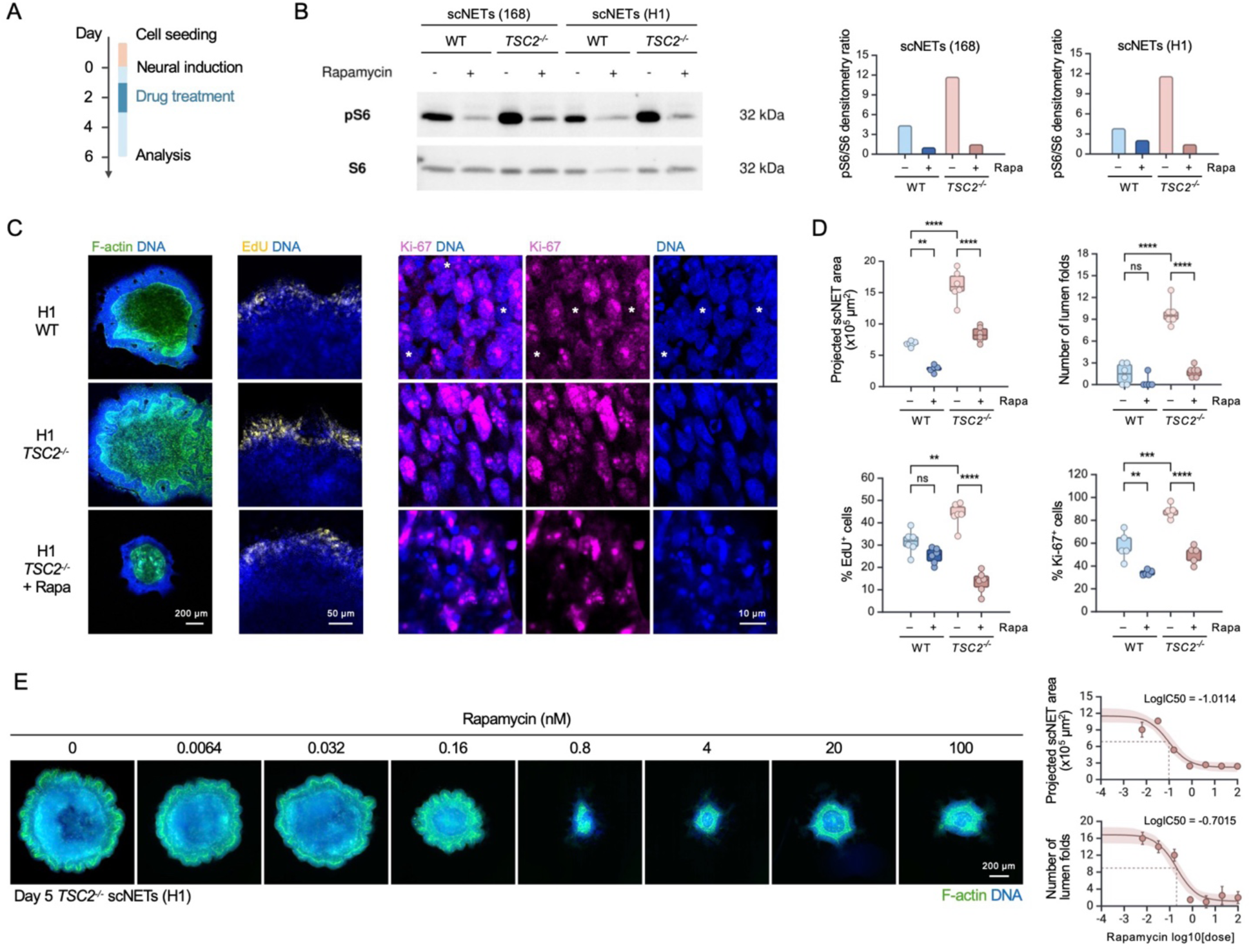
Rapamycin rescues hyperproliferative and folding *TSC2^−/−^* phenotypes. **(A)** Schematic showing the timeline of drug treatment. **(B)** Western blots showing S6 and pS6 levels of the indicated scNETs treated with or without rapamycin. Full blots shown in **Figure S4D**. Boxplots showing the pS6/S6 densitometry ratios. **(C)** Confocal images of day 6 DMSO-treated WT scNET (H1), DMSO-treated *TSC2^−/−^* scNETs (H1), and rapamycin-treated *TSC2^−/−^* scNETs (H1), stained for F-actin (Green), DNA (blue), EdU (yellow), and Ki-67 (magenta). Asterisks mark the cells with no detectable Ki-67 expression. **(D)** Boxplots showing the projected whole area, the number of lumen folds, % of EdU^+^ cells and % of Ki-67^+^ cells of WT scNETs (H1) and *TSC2^−/−^* scNETs (H1), with and without rapamycin. \*\*\*\**p* < 0.00005, \*\*\**p* < 0.0005, ** *p* < 0.005, unpaired *t-*test. **(E)** Fluorescent images showing *TSC2^−/−^* scNETs (H1), treated with rapamycin at indicated concentrations, stained with F-actin (green) and DNA (blue) (left). IC50 curves showing the projected whole pattern area and the number of lumen folds versus the rapamycin dose (right). Scale bars 200 µm, 50 µm and 10 µm are shown. See also **Figure S4**.

**Figure S4.**
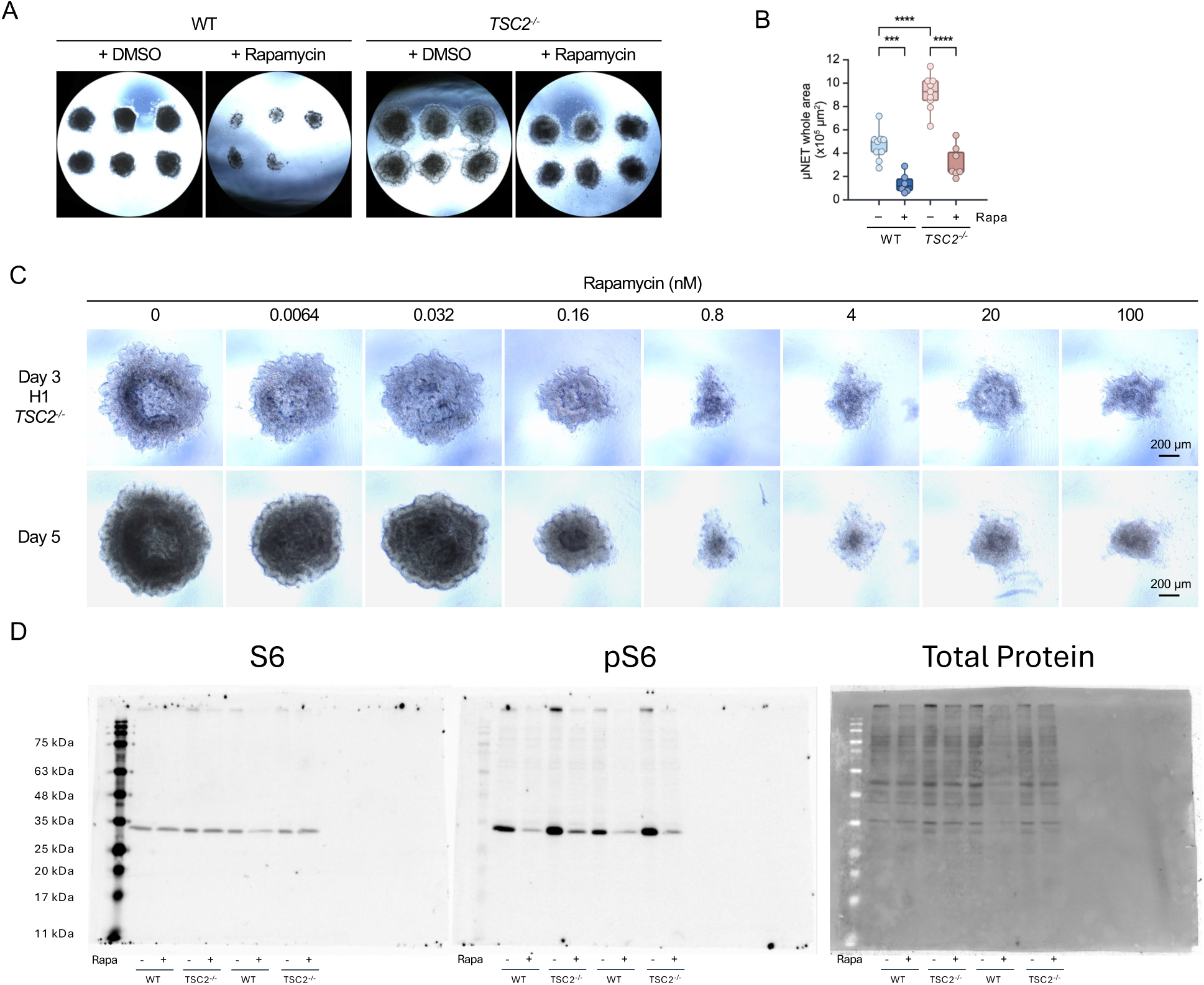
*TSC2^−/−^* scNET phenotypes can be rescued by rapamycin. **(A)** Brightfield images showing scNETs derived from WT hESCs (H1) and those *TSC2^−/−^* hESCs (H1), treated with either DMSO or rapamycin. **(B)** Boxplots showing the projected whole area WT scNETs (168) and *TSC2^−/−^* scNETs (168), treated with and without rapamycin. \*\*\*\**p* < 0.00005, \*\*\**p* < 0.0005, unpaired *t-*test. **(C)** Brightfield images of day 3 and day 5 *TSC2^−/−^* scNETs (H1) treated without or with rapamycin at the indicated concentration. Scale bar indicates 200 µm. **(D)** Full blot data for **Figure 4B**.

To demonstrate the high-content and high-throughput compatibility of our model, we printed ECM droplets in a 96-well plate format and asked if we can determine the rapamycin concentration required to achieve 50% rescue of *TSC2^−/−^* phenotypes, i.e., the IC50 **(Figure 4E)**. *TSC2^−/−^* scNETs were treated with increasing rapamycin concentrations using a 5-fold serial dilution. On day 3, graded responses of *TSC2^−/−^* scNETs to rapamycin in terms of their size and degree of folding were already evident **(Fig S4C)**. Quantitative analyses of the day 5 scNETs revealed rapamycin inhibited tissue size (IC50 = 0.0974 nM) and number of folds (IC50 = 0.198 nM) in a dose-dependent manner **(Figure 4E)**. These IC50 values, being in the nanomolar range, indicated that *TSC2^−/−^* phenotypes were highly sensitive to rapamycin.

In summary, we have reported a versatile bioprinting approach to generate reproducibly sized micropatterned scNETs that show high fidelity to early human brain development. We highlighted the ability of our method to reveal quantifiable disease-associated phenotypes, and its compatibility in high content, high-throughput genetic and chemical compound screens.

## Discussion

In this study, we show that bioprinting can reliably produce droplets of ECM of defined diameter to generate micropatterned scaffolded neuroepithelial tissues (scNETs) when 4% liquid matrigel is added to the media. While this work was done using a commercially available extrusion bioprinter, the rise of open-source, low-cost 3D bioprinting technologies and automated pippetters^44–46^ means that labs will be able to incorporate these models into their workflows more readily. With the conditions described here, we can reliably print ~100 droplets in three minutes and droplet diameter can be controlled by adjusting the print pressure (**Figure S1**). We found that seeding hPSCs onto droplets 700-800 μm in diameter reproducibly generated single lumen scNETs, with neuroepithelial folding occurring in ~ 80% of the tissues. This is in line with previous studies showing that 20% of 500 μm diameter colonies exhibit an open central cavity.^7^ Here, Rho-associated kinase inhibitor (Ri), Y-27632 is added to the media at cell seeding for two hours rather than overnight, because while it is established that Ri promotes survival of dissociated hPSCs,^47^ the ROCK pathway also plays a crucial role in the apical constriction required for neural tube closure.^48^

The standardized and reproducible generation of scNETs with physiologically relevant cell organization make them a unique and accessible tool for studying early neurodevelopment. While many hPSC-derived 2D and 3D *in vitro* neural models exist,^49^ our 6-day protocol can generate neuroepithelial tissues containing self-assembled neural progenitor ventricular zones and even some β-Tubulin^+^ and MAP2^+^ cells in a much shorter time frame and with greater structural fidelity than most existing protocols. This lends the scNET model well to high-throughput drug screening applications. We have demonstrated that matrix droplets can be printed rapidly and in large numbers in a 96 well plate, supporting the utility of our protocol for high-throughput assays. The bioprinting approach also offers greater flexibility than current tools used to produce growth restricted surfaces for micropatterning, as we can easily print in different cell culture vessels and assess the effects of different factors such as ECM substrates, chemical compounds, and culture conditions.

Finally, we provide a unique and powerful tool to quantitatively study the early stages of MCD development. Our observations reflect those seen in cerebral organoids at a later developmental stage, wherein deletion of the tumour suppressor *PTEN*, another inducer of mTORC1-driven MCDs, leads to increased proliferation, cortical size and surface folding.^50^ TSC2-deficiency itself has been shown to alter the structural organization of randomly produced neuroectodermal rosettes in culture and brain organoids;^39,51^ and to cause defects of neural tube closure and morphogenesis in mouse models.^52^ Excess cortical folding is also linked to more severe clinical phenotypes in TS patients.^35^ Our findings contextualize these earlier observations by suggesting the mechanisms underlying morphogenic tissue alterations in MCDs begin at very early developmental stages, supporting hypotheses that neural stem cell projections underlie early stages of cortical folding.^53^ A previous organoid on a chip approach to induce geometrically confined early brain development demonstrated a similar folding phenotype, though this was not prominent until the second week following neuroepithelial induction. This suggests that scNETs may have the potential to capture cortical folding in WT tissues if given more time to develop and that TSC2-deficiency accelerates the normal timeline of tissue folding. To better understand the mechanisms behind cortical folding and other aspects of MCD formation, future work will include dissociation of scNETs from the culture surface to produce free-floating single lumen organoids in extended culture. Additionally, scNETs will be produced with a subpopulation of fluorescently labeled cells to observe cell morphologies and properties of IKNM and ventricular interactions at a single-cell level, and the role of specific proteins that may mediate NPC fate decisions and their structural interactions with the apical lumen.

## Acknowledgments

We thank Dr. William Stanford for generously providing the hPSC lines. We also thank all members of the Julian lab for helpful discussions and technical support, Dr. Lorena Braid’s lab for use of their Incucyte live cell microscope, and Dr. Mahmoud Pouladi for meaningful scientific discussions. This work was supported by funding from the Cancer Research Society (JULIAN, L-CRS 25551), the Natural Sciences and Engineering Research Council of Canada (JULIAN, L-RGPIN-03965) and the New Frontiers in Research Fund (NFRFE-2023-00824). LMJ is a Tier II Canada Research Chair and a Michael Smith Foundation for Health Research/Parkinson Society BC Scholar. LL is funded by MITACS and the NSERC Alliance program and SN by SFU and Phyllis Carter Burr Graduate Fellowships.

## Author contributions

Conceptualization: N.I.F, K.K.L.W, G.A., L.M.J; Validation: K.K.L.W, G.A.; Methodology and Resources: N.I.F., K.K.L.W, G.A., A.M., L.L., S.N.; Investigation: K.K.L.W, N.I.F, A.M; Writing (Original draft & review): K.K.L.W, N.I.F, L.M.J.; Supervision: L.M.J.; Funding acquisition: L.M.J.

## Method details

### Cell models

Human PSC lines used in this study were 168 (iPSC, male) and H1 (hESC, male; WiCell WA01). Control 168 line was generated previously and reported in^54^. Isogenic *TSC2^−^*^/−^ 168 and H1 lines were generated previously.^16^ All cell lines tested negative for mycoplasma.

### Cell culture

Human PS cell lines were maintained in a standard feeder-free culture system using mTeSR medium (STEMCELL Technologies) on Matrigel hESC-Qualified Matrix (Corning 354277), cultured in 6-well tissue culture plates. Cell culture was visually examined during each passage to ensure the absence of spontaneously differentiated, mesenchymal-like cells in culture.

### 3D printing of extracellular matrix

Extracellular matrix (ECM) droplets were printed using a CELLINK BIO X6 bioprinter onto Nunc Thermanox coverslips (Thermo Scientific #174969) in a 24-well tissue culture plate. Empty wells in the plates were filled with sterile water, creating a humid environment to prevent droplet evaporation. Cultrex (R&D Systems #BME001-10) prepared at 1.5x concentration was added into a BIO X6 print cartridge. A 27-gauge needle was then fitted to the print cartridge along with the air adapter connector. The assembled cartridge was then inserted into the print position of the printer. The BIO X6 printer software used was DNA studio 3 with the print protocol setup using the Model function directed by G-code script. Print pressure was set to 15 kPa, unless otherwise indicated. To minimize droplet evaporation, print bed temperature was set to 14°C and the internal fan was disabled. After print head calibration, the print job was released. Once ECM printing was complete, the plate was sealed with parafilm and refrigerated for at least 2 hours before cell seeding. The vessels with printed droplets were stored at 4°C for up to 2 weeks before use. Droplet size was controlled with the g-code script at a diameter of 800 µm. Before cell seeding, the wells with printed ECM droplets were spray washed with PBS. The G-code scripts used to generate micropatterns have been deposited in Open Science Framework https://osf.io/538qa/.

### Generation of scNETs

Colonies of human PS cells grown to a confluence of >70% were dissociated using Accutase (STEMCELL Technologies #07920) at 37°C for 6 min. Single cells were resuspended in mTeSR supplemented with 10 µM ROCK inhibitor Y-27632 (STEMCELL Technologies), followed by seeding at the indicated density onto the Cultrex-coated coverslips. After 2 hours, the ROCK inhibitor was removed by replacing the media with fresh mTeSR. Once the hPSCs fully covered the printed droplet area, neural induction and lumenogensis (Day 0) was initiated. Cells were treated with neural induction media (NIM) containing DMEM/F12 (Gibco, 11320-033), Neurobasal (Gibco 11320-033), 1% N2 (Gibco, 17502-048), 2% B27 (Gibco, 17504-044), 1x Glutamax (Gibco 35050-061), 1% MEM (Gibco, 11140-050) with dual SMAD inhibitors SB-431542 (Stemcell Technologies, 72234, 10 µM) and LDN (Stemcell Technologies, 72147, 500nM) with 4% matrigel. Full media was done every 2 days post-neural induction using NIM with 2% matrigel. For rapamycin treatment, the drug was added to the cells on Day 1 for 48 hours at 20 nM, unless otherwise indicated.

### Immunostaining of scNETs

Cells were rinsed in PBS and fixed in 4% PFA in PBS for 15 min at room temperature. The cells were then washed with PBS with 0.1% Triton X-100 (PBST), followed by incubation with 5% normal goat serum (NGS) in PBST for 1 hour at room temperature. After blocking, the cells were incubated with primary antibodies overnight at 4°C. Primary antibodies used include rabbit anti-PAX6 (1:1000; Proteintech #12323-1-AP), rabbit anti-Ki-67 (1:200; Proteintech #27309-1-AP), mouse anti-ZO-1 (1:200; Invitrogen #33-9100), mouse anti-β-Tubulin III (1:500; STEMCELL Technologies #60100), mouse anti-N-cad (1:500; Cell Signalling #14215), chicken anti-MAP2 (1:5000; Millipore Sigma #AB5543). After washing with PBST, the cells were incubated with secondary antibodies, phalloidin and DAPI in the dark overnight at 4°C, followed by a final wash, mounting and imaging.

### Image acquisition and processing

Images were taken on a laser scanning microscope (Zeiss LSM880 with Airyscan) and processed by Image J. Time-lapse imaging was performed on the Incucyte Live-Cell Analysis System (Sartorius).

### EdU assay

EdU assay was performed using Click-iT™ EdU Cell Proliferation Kit for Imaging, Alexa Fluor™ 488 dye (Invitrogen #C10337). Briefly, scNETs were treated with 10 µM of EdU for 2 hours, followed by fixation and PBST wash. Click chemistry was performed by incubating the fixed samples with the Click-iT reaction mix cocktail for 30 minutes at room temperature in the dark. After PBST wash, the samples were blocked with NGS for downstream immunostaining.

### RT-qPCR

Total RNA isolation was performed using Quick-DNA/RNA Microprep Plus Kit (Zymo Research D7005), followed by cDNA synthesis using High-Capacity cDNA Reverse Transcription Kit (Applied Biosystems 4368814). RT-qPCR was performed using PowerUp™ SYBR™ Green Master Mix (Applied Biosystems A25742) on CFX384 Real-Time System (Bio-Rad). See **Table S1** for primers used.

### PCR

*TSC2* expression was knocked out by introduction of a frame-shift 35-base stop codon sequence at *TSC2* exon 3, which is confirmed by PmeI digestion. Genomic DNA isolation was performed using the Quick-DNA/RNA Microprep Plus Kit (Zymo Research D7005). PCR reaction was performed using OneTaq Quick-load 2x Master Mix (New England Biolabs M0486) with 300 ng of isolated template DNA and TSC2 primers with cycling parameters of: 94°C for 30 sec, 25x [94°C 20s, 56.5°C 30 sec, 68°C 30s], 68°C for 5 min, 4°C for ∞. The PCR product then underwent a restriction digest reaction with PmeI (New England Biolabs R0560) in CutSmart Buffer (New England Biolabs B6004S) for 1 hr at 37°C, followed by heat inactivation at 65°C for 20 min. Samples were loaded in 10x BlueJuice Gel Loading Buffer (Thermo Fisher 10816015) and run for 30 minutes at 100V on a 1% agarose gel and imaged using ethidium bromide on the ChemiDoc MP Imaging System (Bio-Rad). See **Table S1** for primers used.

### Western blotting

Cells were lysed with 1x RIPA Buffer (Sigma-Aldrich R0278), supplemented with Pierce Protease Inhibitor (ThermoFisher A32963) and Phosphatase Inhibitor Cocktail (Millipore Sigma 524629). Protein lysates with 1X SDS sample buffer were resolved on SDS/PAGE and then transferred to nitrocellulose membranes. Membranes were blocked with 5% bovine serum albumin for 1 hour before overnight primary antibody incubation. Subsequently, the blots were incubated in fluorescent secondary antibodies for 2 hours. Blots were then imaged on the ChemiDoc MP Imaging System (Bio-Rad) and target protein was quantified using Image Lab Software (Bio-Rad). See **Key Resources Table** for antibodies used.

### Quantification and statistical analysis

Data were presented by mean ± SEM obtained from at least two biological replicates. *p* values were determined by unpaired t tests. *p* < 0.05 was considered statistically significant.

## Video Legends

**Video 1.** Time-lapse imaging of micropatterned scNET development from day 0 to day 6.

**Video 2.** 3D visualization of day 6 micropatterned H1 wildtype scNET **(A)** and that of H1 *TSC2^−/−^* scNET **(B)**.

**Video 3.** Time-lapse imaging of micropatterned *TSC2^−/−^* scNET development from day 1 to day 6.

