## Supplementary material for "A bioprinting approach for high-throughput production of micropatterned neuroepithelial tissues and modeling TSC2-deficient brain malformations": Table S1

**Table S1.** **Primer sequences used in qPCR and PCR**

| Oligonucleotides | | |
|---|---|---|
| qPCR primers for TBP<br>Forward: 5'-CAAGAACTTAGCTGGAAAACCC-3'<br>Reverse: 5'-GATAAGAGAGCCACGAACCAC-3' | Integrated DNA Technologies | Hs.PT.39a.22214825 |
| PCR primers for TSC2<br>Forward: 5’-TCC TCG GGA TGG AGC AGT AA-3’<br>Reverse: 5’-TGC AAA CCA GAT CAT CGG CA-3’ | *Delaney et al., 2020* | |
| qPCR primers for PAX6<br>Forward: 5'-GACACCACCGAGCTGATTC-3'<br>Reverse: 5'-ATTTCAGAGCCCCATATTCGAG-3' | Integrated DNA Technologies | Hs.PT.58.3002797 |
| qPCR primers for MAP2<br>Forward: 5’-ATCTTGACATTACCACCTCCAG-3'<br>Reverse: 5'-TGAAGAACATCCGCCACAG-3' | Integrated DNA Technologies | Hs.PT.58.1947612 |
| qPCR primers for FOXG1<br>Forward: 5'-CGTCCACCATATAGTTCCATGA-3'<br>Reverse: 5'-TGACTGCTTTGCCATTTCATTC-3' | Integrated DNA Technologies | Hs.PT.58.26906112.g |
| qPCR primers for NKX2.1<br>Forward: 5'-TGCCGCTCATGTTCATGC-3'<br>Reverse: 5'-CAGGACACCATGAGGAACAG-3' | Integrated DNA Technologies | Hs.PT.58.2461055 |
| qPCR primers for TUBB3<br>Forward: 5'-GGCCTTTGGACATCTCTTCAG-3'<br>Reverse: 5'-CCTCCGTGTAGTGACCCTT-3' | Integrated DNA Technologies | Hs.PT.58.20385221 |
| qPCR primers for VIM<br>Forward: 5'-CAAGACCTGCTCAATGTTAAGATG-3'<br>Reverse: 5'-GTGAATCCAGATTAGTTTCCCTCA-3' | Integrated DNA Technologies | Hs.PT.58.38906895 |
